# Opening the black box of high resolution fish tracking using yaps

**DOI:** 10.1101/2019.12.16.877688

**Authors:** Henrik Baktoft, Karl Ø. Gjelland, Finn Økland, Jennifer S. Rehage, Jonathan R. Rodemann, Rolando Santos Corujo, Natasha Viadero, Uffe H. Thygesen

## Abstract

The R package `yaps` was introduced in 2017 as a transparent open source alternative to closed source manufacturer-provided solutions to estimate positions of fish (and other aquatic animals) tagged with acoustic transmitters.

Although `yaps` is open source and transparent, the process from raw detections to final tracks has proved to be challenging for many potential users, effectively preventing most users from accessing the benefits of using `yaps`. Especially, the very important process of synchronizing the hydrophone arrays have proven to be an obstacle for many potential users.

To make `yaps` more approachable to the wider fish tracking community, we have developed and added user-friendly functions assisting users in the complex process of synchronizing the data.

Here, we introduce these functions and a six-step protocol intended to provide users with an example workflow that can be used as a template enabling users to apply `yaps` to their own data. Using example data collected by an array of Vemco VR2 hydrophones, the protocol walks the user through the entire process from raw data to final tracks. Example data sets and complete code for reproducing results are provided.

## Introduction

Tracking of free-ranging fish in the wild with high spatiotemporal resolution using acoustic transmitters and fixed position hydrophone arrays is an increasingly popular methodology to obtain otherwise intractable information about fish behaviour and ecology (Lennox et al. 2017). Until recently, the software needed to estimate positions from detections was available only through manufacturer-supplied solutions with little to no transparency, leaving users (i.e. scientists and resource managers) in the blind regarding which algorithms, filters, etc. had been applied to the data. The R package `yaps` (Yet Another Positioning Solver) was introduced in 2017 as an alternative to these proprietary and vendor-specific options for estimating positions and tracks (Baktoft et al. 2017). Although `yaps` was developed with an intent of openness, transparency, reproducibility and flexibility, `yaps` is not a turnkey or a one-size-fits-all solution. While `yaps` itself is open and transparent, the process from collecting raw data in the field to having final estimated tracks is long and sometimes challenging. Especially, the very important process of synchronizing the hydrophone arrays has proven to be a major obstacle for many potential users.

### Synchronization of hydrophone arrays

Regardless of the algorithm used for positioning, synchronization of the hydrophone array prior to track estimation is extremely important for quality of the final tracks and great care should be taken to achieve the best possible synchronization. The newly developed synchronization method recently included in the `yaps` package has several benefits. It synchronizes all hydrophones in one go, avoiding sequential propagating synchronization, which may lead to error accumulation in the perimeter of the hydrophone array. Additionally, it allows for estimation of hydrophone positions, which can be useful if the initial hydrophone positions are uncertain e.g. due to deep water, strong current or use of a low-accuracy GPS. Perhaps most importantly, synchronization in `yaps` is aimed at being user-friendly. To meet this, we have attempted to wrap the necessary complex mathematical operations and data wrangling inside easy-to-use functions. Hydrophone internal clocks drift in varying directions and the rate of drift can be affected by external factors such as water temperature. Therefore, temporally flexible non-linear correction is needed. This is implemented in the `yaps` synchronization procedure through multiple hydrophone-specific polynomials.

### Motivations for developing and using yaps

- **Transparency and reproducibility**. These concepts should be fundamental in all science and scientific methodology. However, available proprietary software for estimating positions lack transparency and constitute “black boxes” making reproducibility and scrutiny of results very challenging, if not impossible.
- **Open source**. In addition to complete transparency, publishing `yaps` open-source enables the user community to implement study specific adaptations, tweak parts of the estimation models and contribute to the continued development. We urge users to submit improvements, refinements, etc. to the main `yaps` repository on github, thereby helping future users.
- **Vendor agnostic**. The basic input to `yaps` is plain matrices of time-of-arrivals of transmitter signals detected by hydrophones. The brand of hydrophones and transmitters are irrelevant. This enables more direct comparisons of tracks and of the derived behavioural quantifications among studies using varying vendors.
- **Get the most out of your data**. To our knowledge, all vendor supplied tracking software processes data from individual pings independently from other pings. By doing this, these algorithms explicitly discard collected data when pings are detected by less than three hydrophones. However, these data are valuable for estimating tracks if used correctly, and can dramatically increase overall data yield from your study (see (Baktoft et al. 2017)). Note, that this otherwise discarded data is only of value if the transmitter burst interval is either fixed or the sequence of random burst intervals is known.
- **Get the best out of your data**. The holistic complete-track-oriented approach used by `yaps` has proven to be superior in many instances to the isolated single-ping-oriented approach used in vendor software. The single-ping approach is prone to numerical challenges, which can lead to varying degrees of spurious outliers, multiple position estimates from individual pings (sometimes known as “ghost positions”) and overall increased track jaggedness and uncertainties. These artifacts often necessitate the use of post-processing filters and/or smoothing techniques. In contrast, many of these irregularities are, in most circumstances, handled well by `yaps` and resulting tracks are often ready for use in further analyses. Note, we are not stating that `yaps` estimated tracks never need or will benefit from post-processing – it depends very much on study specifics such as the acoustic environment and transmitter burst interval type and duration. However, using `yaps`-estimated tracks as the starting point for post-processing has in all cases we know of, been superior to using tracks estimated by vendor software.

## Methods and Materials

### A six step protocol from raw detections to synchronized data

The workflow going from raw detection to synchronized data is a multi-step process. Below we describe each step in detail, and then provide the concise code to implement each step.

1. **Prepare data** as a list containing tables `hydros`, `detections` and optionally `gps` following the format in the included example data set `ssu1` described below. The table `gps` is optional and only relevant for test tracks. To aid in generating table `detections` from hydrophone downloads, we have included a helper function `prepDetections()` in `yaps`.
2. **Set synchronization parameters**. Each data set is different in terms of hydrophone array configuration, precision and accuracy of hydrophone positioning, acoustic environment and detection probability, synchronization tag configuration, manufacturer etc. A number of parameters are available in function `getInpSync()` to setup the synchronization process for best results.
  - `max_epo_diff` For the synchronization process to function properly, hydrophones are assumed to be roughly synchronized initially (e.g. by setting correct time via pc communication), such that trains of emitted signals from synchronization transmitters can be aligned correctly across all hydrophones. `max_epo_diff` sets the upper threshold for differences in time of arrival (TOA) of a synchronization transmitter signal detected at each hydrophone. Best choice of parameter value depends on burst rate of synchronization transmitter and how far apart the internal clocks of the hydrophones are prior to synchronization. This parameter value should never exceed half of minimum synchronization transmitter burst rate. A linear time correction applied to the detection data prior to synchronization can dramatically improve the alignment process.
  - `min_hydros` To ensure connectivity throughout the array, pings from synchronization transmitters need to be detected by multiple hydrophones. Additionally, every hydrophone need to detect synchronization transmitters also detected by one or more other hydrophones. Therefore, positioning of synchronization transmitters should be done in such a way that multiple hydrophones can recieve it’s pings and such that all hydrophones can recieve pings from at least one synchronization transmitter also detected by one or more other hydrophones. `min_hydros` sets the lower threshold of number of hydrophones needed to detect each synchronization transmitter ping in order to be included in the synchronization process. This value should be as high as possible, while ensuring that all hydrophones are included. If `min_hydros` is too high, isolated hydrophones risk completely falling out of the synchronization process. Future versions will work towards automation of this step.
  - `time_keeper_idx` One hydrophone is assigned the role of time-keeper. All other hydrophones are synchronized to match the time-keeper. `time_keeper_idx` is the index of this hydrophone specified in the `hydros` table (for instance, the hydrophone with smallest overall clock-drift).
  - `fixed_hydros_idx` Vector of hydrophone indexes specified in the table `hydros` of hydrophones where the position is assumed to be known with adequate accuracy and precision. As many as possible fixed hydrophones should be included in order to reduce overall computation time and reduce variability. As a bare minimum two hydrophones need to be fixed, but we strongly advice to use more. If the position determined by gps of one or more hydrophones is uncertain (e.g. placed at deep water) or missing all together, their position can be estimated during the synchronization process, by excluding their index from this vector.
  - `n_offset_day` Specifies the number of hydrophone specific polynomials to use per day. For PPM-based systems like Vemco VR2 and Thelma TBR700, 1 or 2 is often as good choice.
  - `n_ss_day` Specifies number of speed of sound estimates per day. Future versions will enable use of logged water temperature instead. However, estimating speed of sound yields an extra option for quality checking the final synchronization model.
  - `keep_rate` Synchronizing large data sets can take a long time. However, there is often excess number of synchronization transmitter detections available and a sub-sample is typically enough to obtain a good synchronization model. This parameter specifies the proportion (0-1) of the data to keep when sub-sampling. Future versions will work towards automation of this step.
3. **Compile input data** for the synchronization process using function `getInpSync()`.
4. **Obtain synchronization model** by running function `getSyncModel()`. This can take a long time for larger data sets including more hydrophones and/or spanning longer periods. In such cases, we suggest to start with the first few days to make sure everything looks reasonable before attempting to synchronize the entire data set. Also consider using the parameter `keep_rate` in `getInpSync()`.
5. **Quality check the synchronization model** to ensure the hydrophone array is synchronized well. Functions `plotSyncModelResids()` and `plotSyncModelCheck()` produce plots which can be used to diagnose if the overall model fit is good.
  - Function `plotSyncModelResids()` plots synchronization model residuals (in meter). Residuals centered closely around zero indicate that the synchronization model is good. If fixed hydrophones and/or synchronization transmitters consistently have large deviations from zero, it may indicate that initial position accuracy of the hydrophone is sub-optimal. Either obtain a better position (typically not possible) or allow estimation of the position of that particular hydrophone (i.e. remove it’s idx from vector `fixed_hydros_idx`). Note that it requires a rerun of the entire process for altered synchronization parameters to take effect.
  - Function `plotSyncModelCheck()` applies the synchronization model to the data obtained by `getInpSync()` and compares true distances between hydrophone and synchronization transmitters to distances estimated based on the newly synchronized TOA-data. Similar to `plotSyncModelResids()`, consistent large deviations indicate a problem in the synchronization model that needs further attention.
6. **Apply the synchronization model** to the entire data set using `applySync()`.

The data should now be synchronized and ready to be processed by `yaps` to obtain estimated tracks.

### Package installation and dependencies

`yaps` is an R package available from a github repository at github.com/baktoft/yaps. To install in R, package `devtools` (Wickham, Hester, and Chang 2019) is needed. Integral parts of `yaps` relies on package `TMB`, Template Model Builder (Kristensen et al. 2016). Run the following lines in R to meet dependencies, ensure `TMB` is working and install `yaps`.

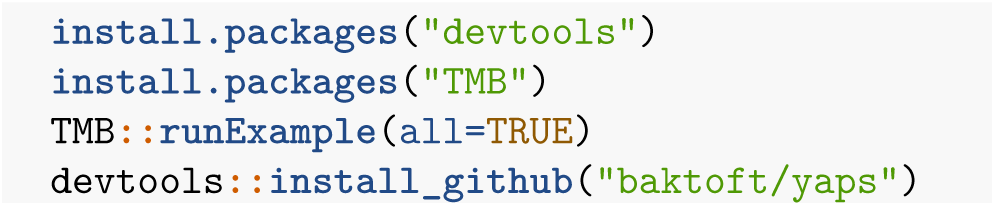

Note that `Rtools` might also be needed. If so, go to https://cran.r-project.org/bin/windows/Rtools and choose the correct version for your current installed version of R. These steps are only needed once.

### Example data in package yaps

We use the `ssu1` data set included in `yaps` for the purpose of this demonstration. This is a small data set containing test tracks collected in Florida Bay, Florida, USA using 19 Vemco VR2 hydrophones. The size of the data set makes it suitable for demonstration purposes. The data set is a `list` containing the following three `data.tables`: `hydros`, `detections` and `gps`. The table `hydros` contains positions of the hydrophones obtained using a handheld gps-unit, serial number of co-located synchronization transmitters and hydrophone index (must be in range `1:nrow(hydros)`); table `detections` contain all detections of synchronization transmitters and transmitters used for the test track; and table `gps` contains gps-recordings of the test track used as ground truth when evaluating quality of the estimated tracks. See `ssu1` help (type `?ssu1` into R console) for further information.

### Applying the six step protocol to synchronize example data ssu1

#### 0. Load the yaps package

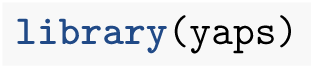

#### 1. Prepare data

We can use the function `prepDetections()` to generate the `data.table detections` from raw detections download from all hydrophones in the array. Future versions will work towards support for other vendors and data formats. The included example data set `ssu1` is a subset of the `data.table vue` created below.

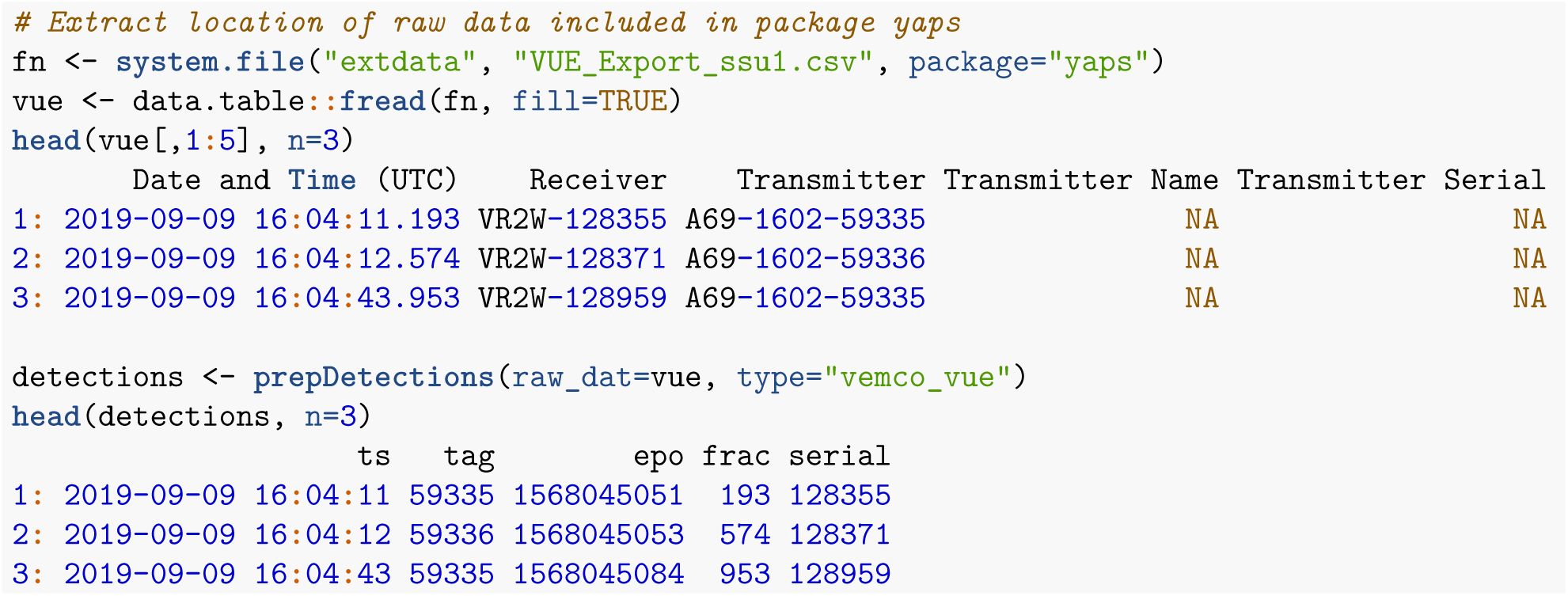

#### 2. Define synchronization parameters

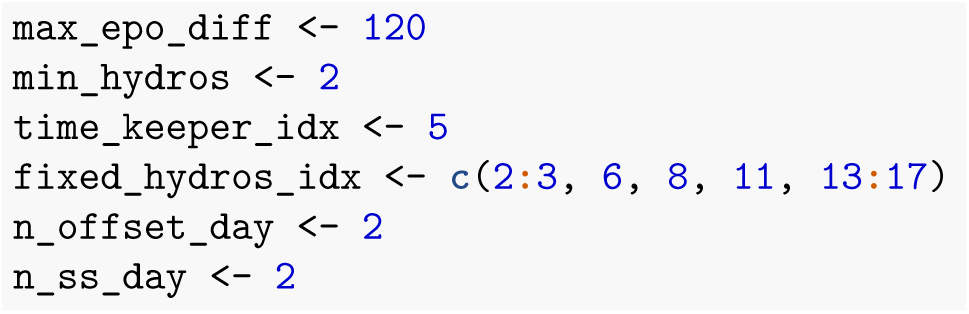

#### 3. Compile input data

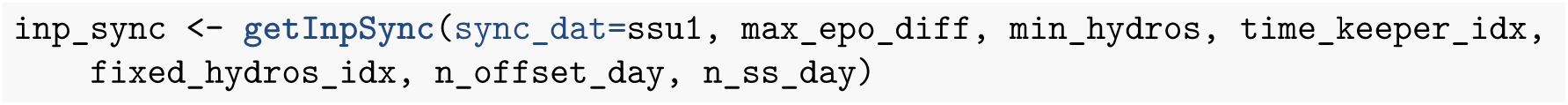

#### 4. Obtain synchronization model

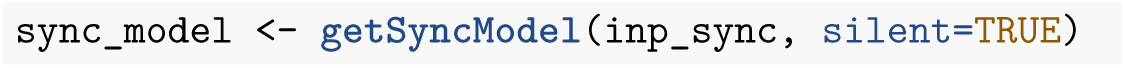

#### 5. Quality check the synchronization model

by plotting model residuals (Fig. 1, 2 & 3) and model check plots (Fig. 4, 5, 6 & 7). For all plots, values closer to zero are better.

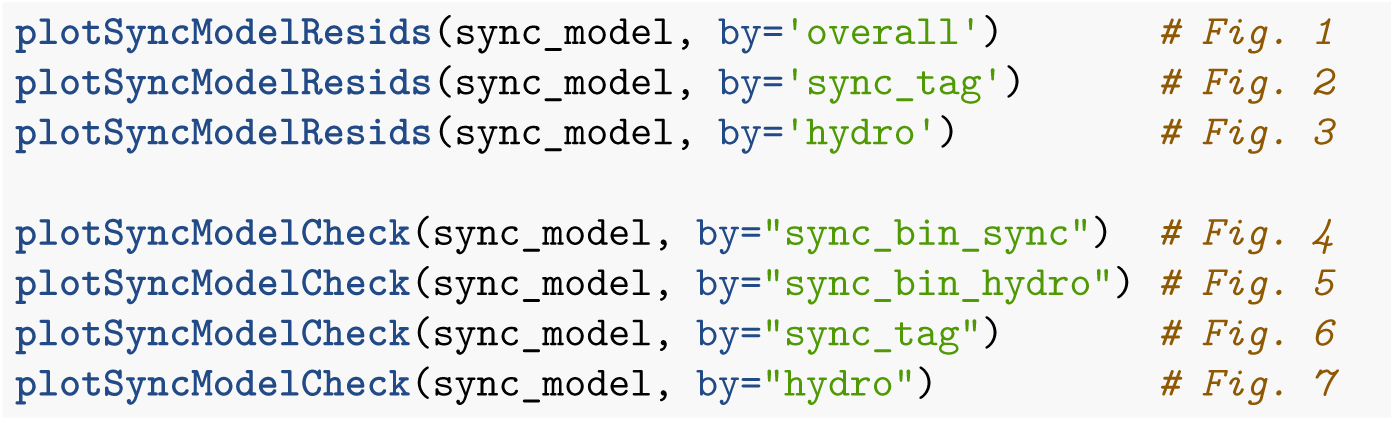

**Figure 1:**
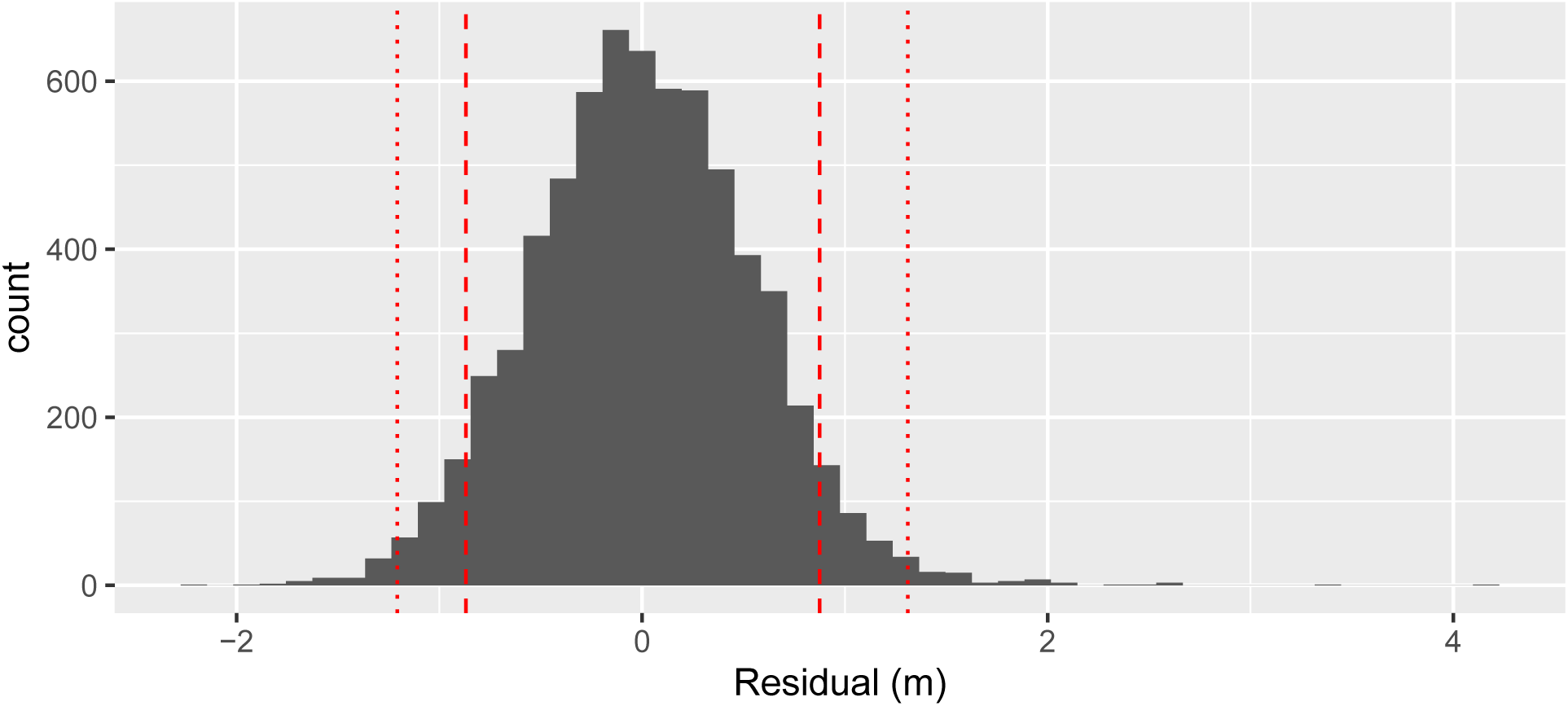
Overall histogram of sync model residuals (converted to meter). Vertical red lines indicate 1, 5, 95 and 99 % quantiles.

**Figure 2:**
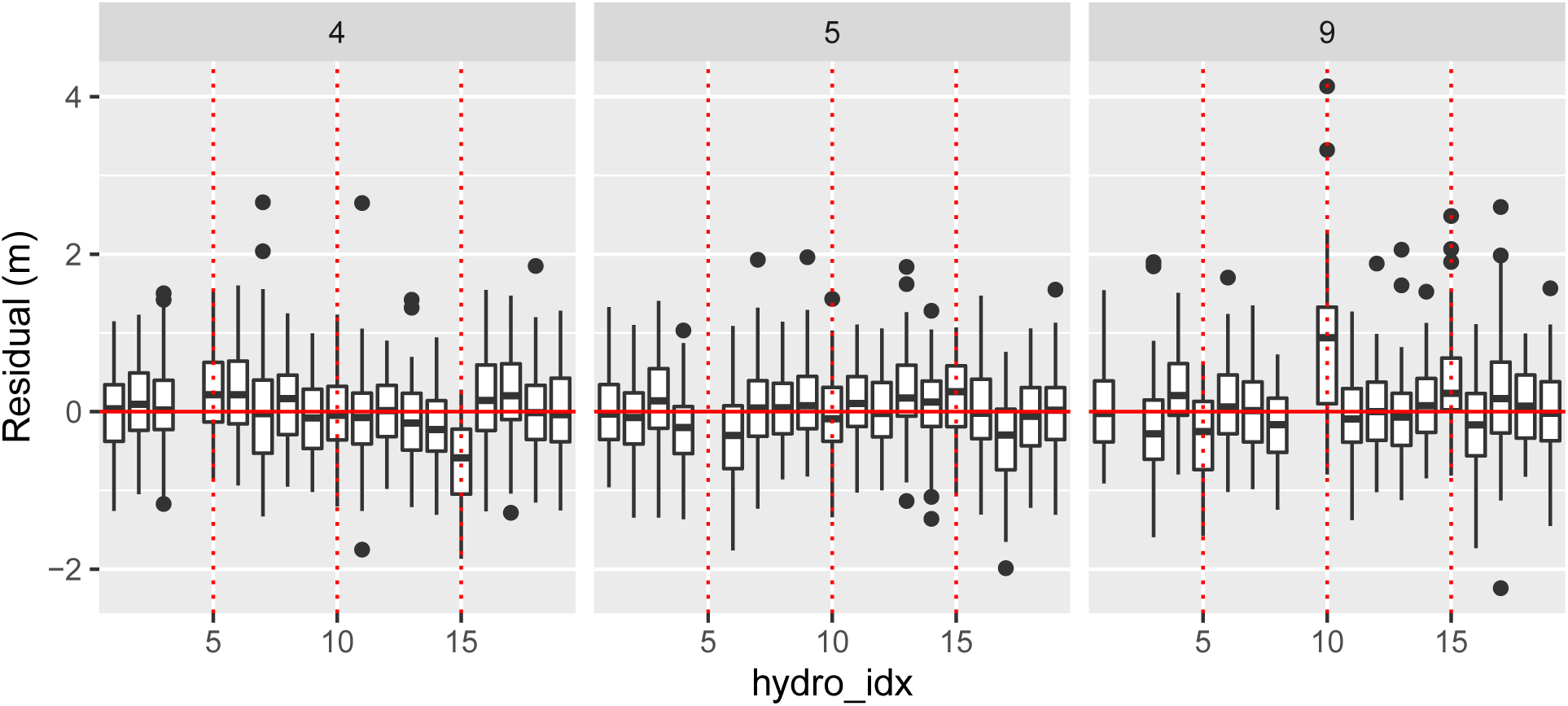
Boxplots of sync model residuals (in meter) grouped by sync tags (panel) and hydrophone (x-axis). Vertical red lines added for every fifth hydro_idx to aid in identifying troublesome hydrophones

**Figure 3:**
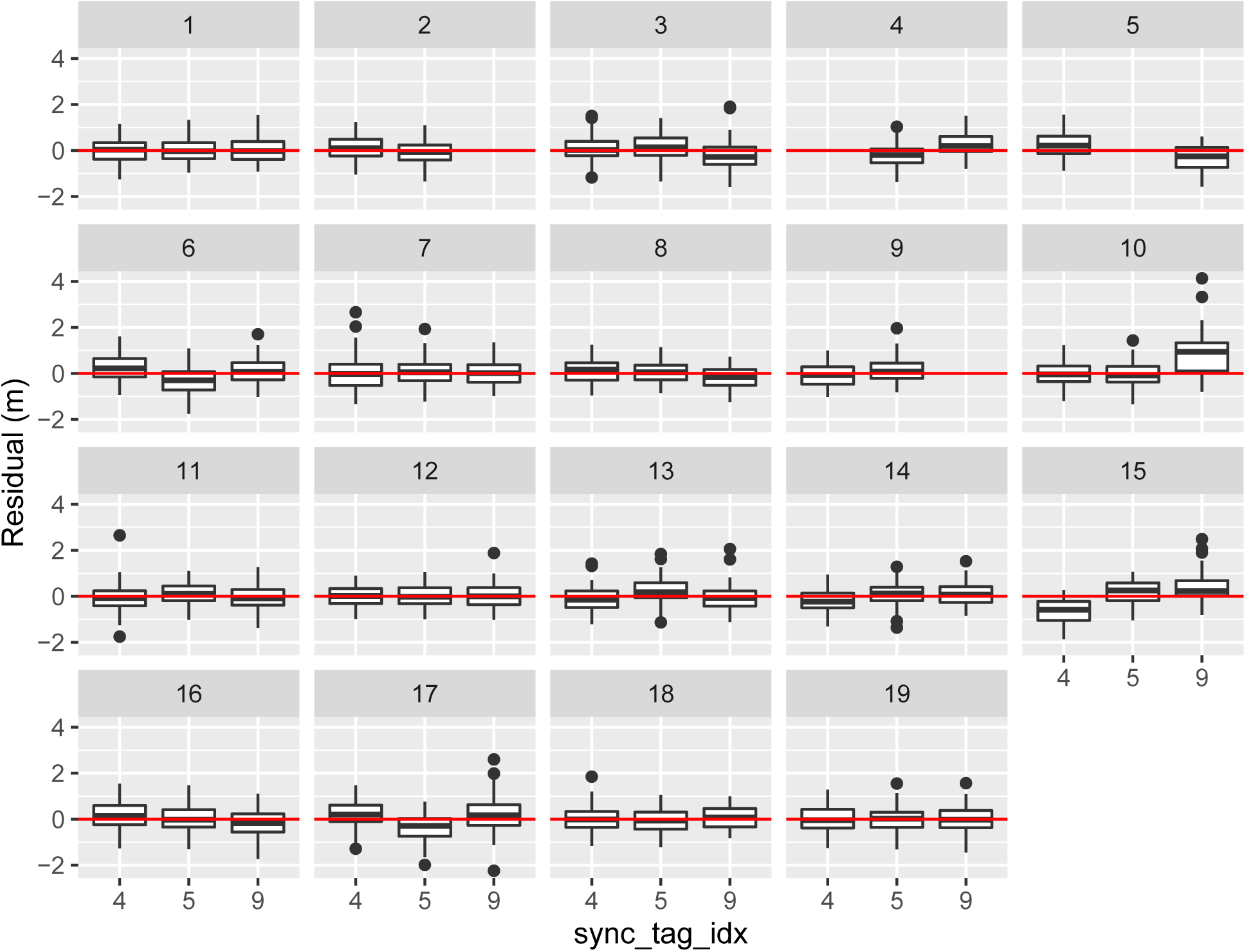
Boxplots of sync model residuals (in meter) grouped by sync tags (x-axis) and hydrophone (panels)

**Figure 4:**
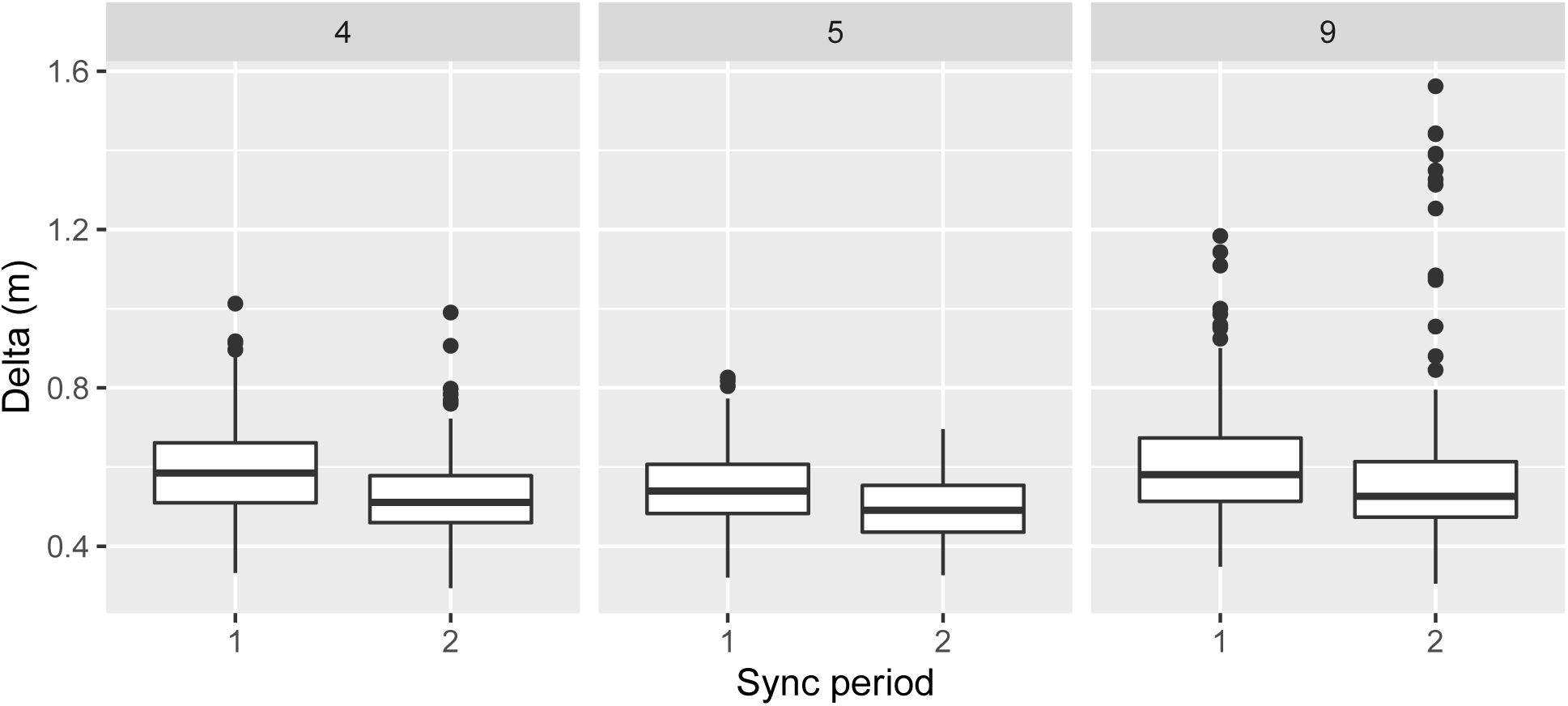
Delta grouped by sync tag (panel) and sync period (x-axis)

**Figure 5:**
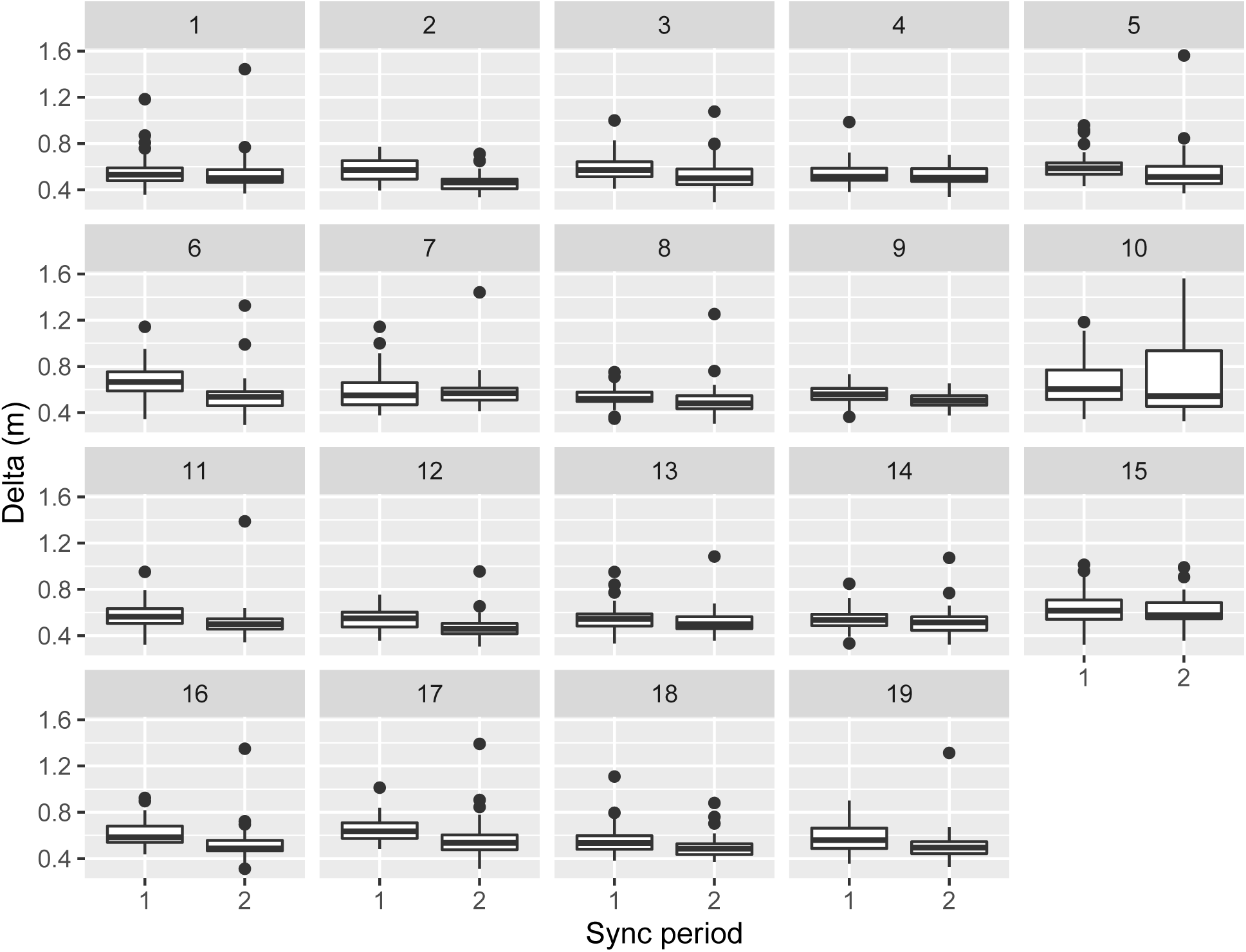
Delta grouped by hydro_idx (panel) and sync period (x-axis)

**Figure 6:**
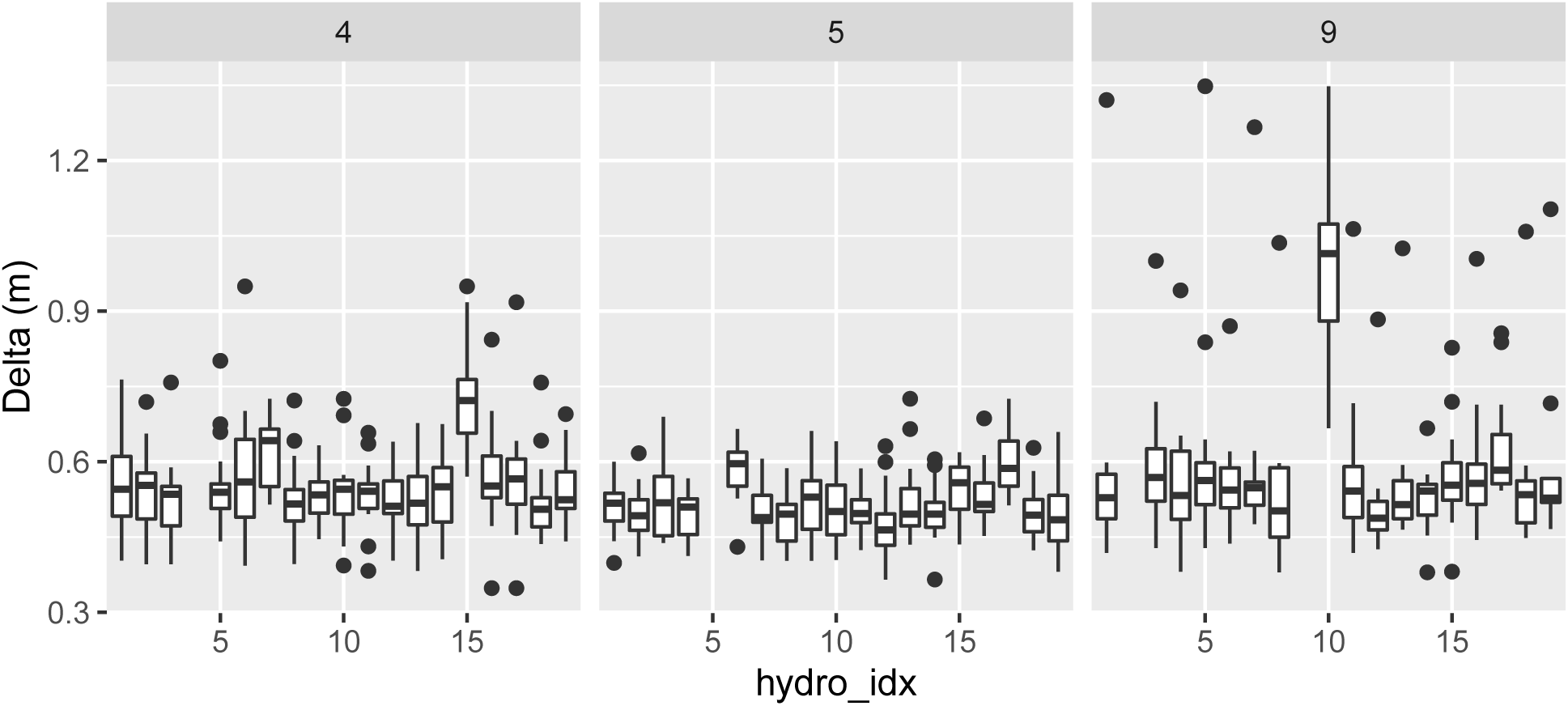
Delta grouped by sync tag (panel) and hydro_idx (x-axis)

**Figure 7:**
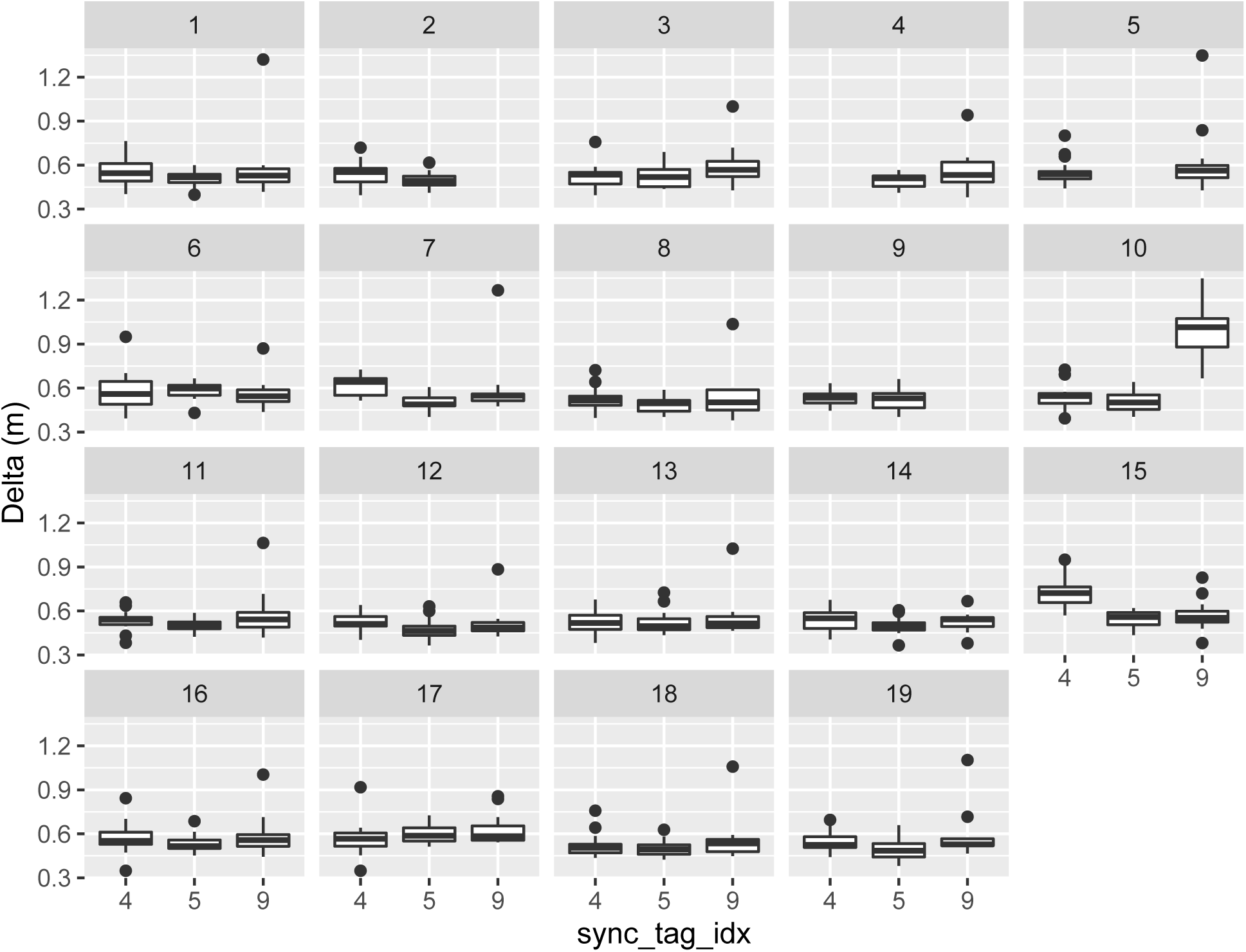
Delta grouped by hydro_idx (panel) and sync tag (x-axis)

#### 6. Apply the synchronization model

to all data in table `detections`. Synchronized timestamps are found in column `eposync`. Note, fractions of seconds are not shown in output below. Data are now synchronized and ready to track estimation using `yaps`.

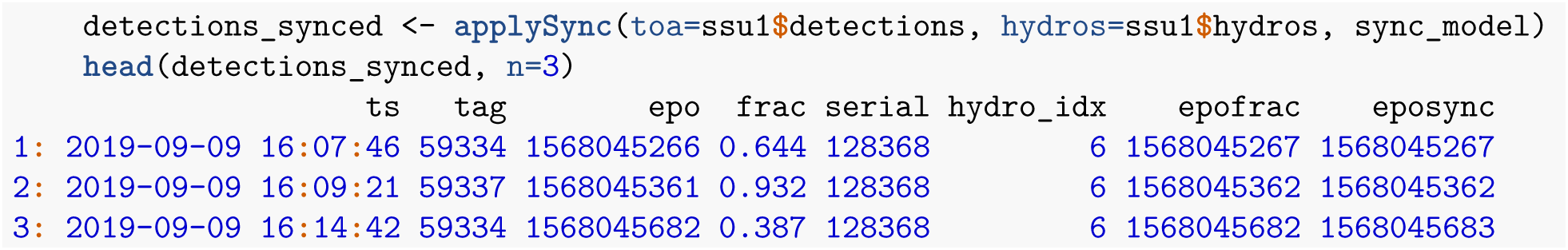

### Estimate track using yaps on newly synchronized ssu1 data

The only tracks to estimate from the `ssu1` data set are test tracks performed to test feasibility of deploying an array at this specific location. In the original data set, several transmitters with different power and burst rates were used. Here, we focus on a single of these: ID 15266, Vemco V9, high power output, 20-40 second burst interval.

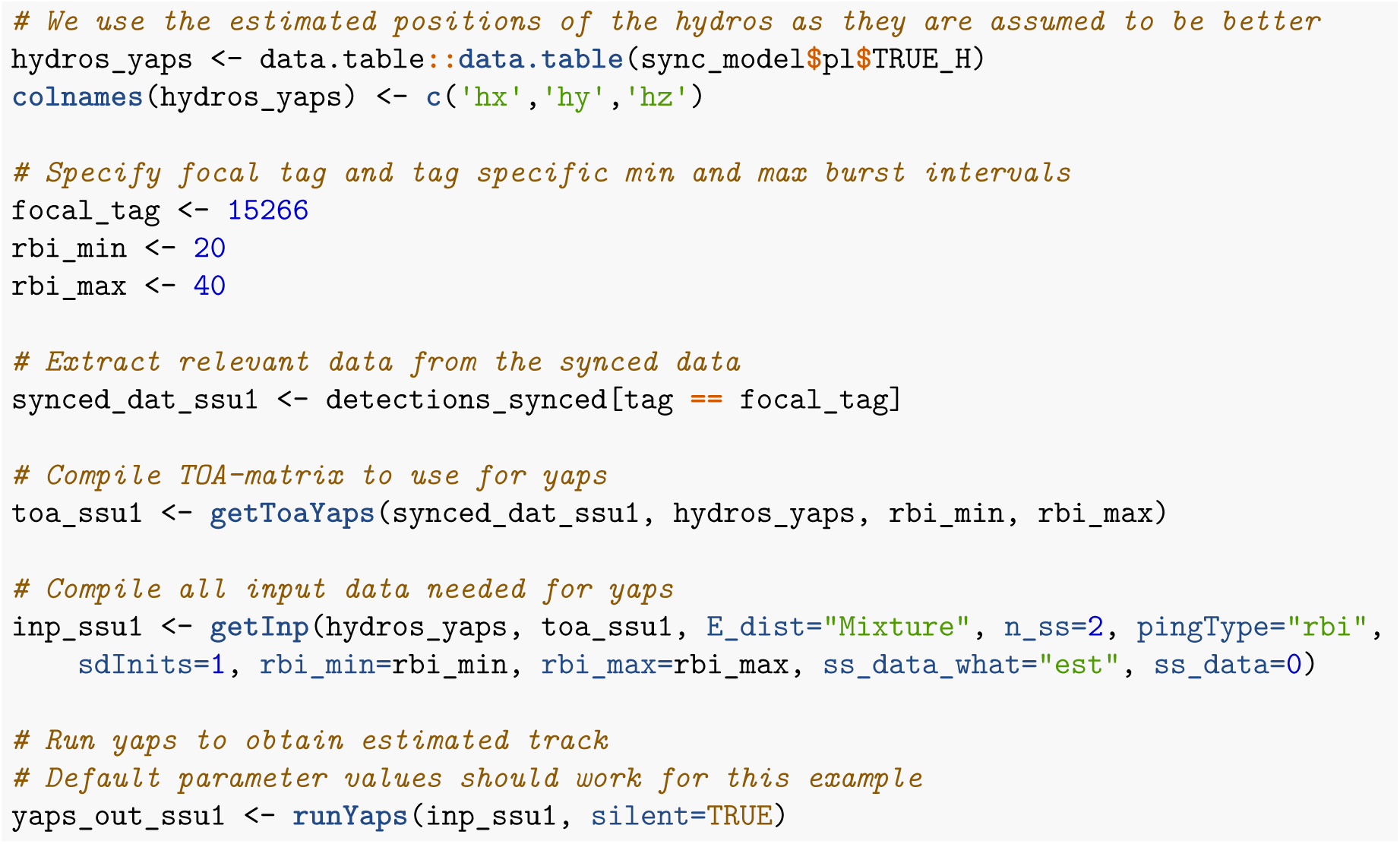

## Results

### Validating the estimated track - ssu1

The code below produces a basic plot of `yaps`-estimated track after applying the six-step synchronization protocol.

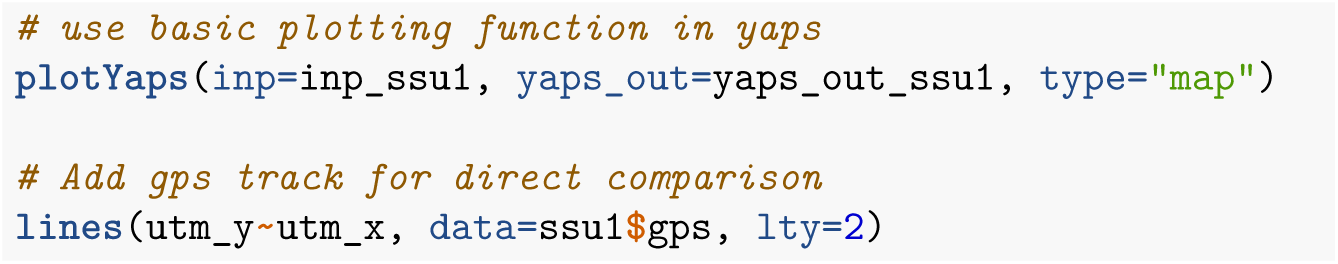

**Figure 8:**
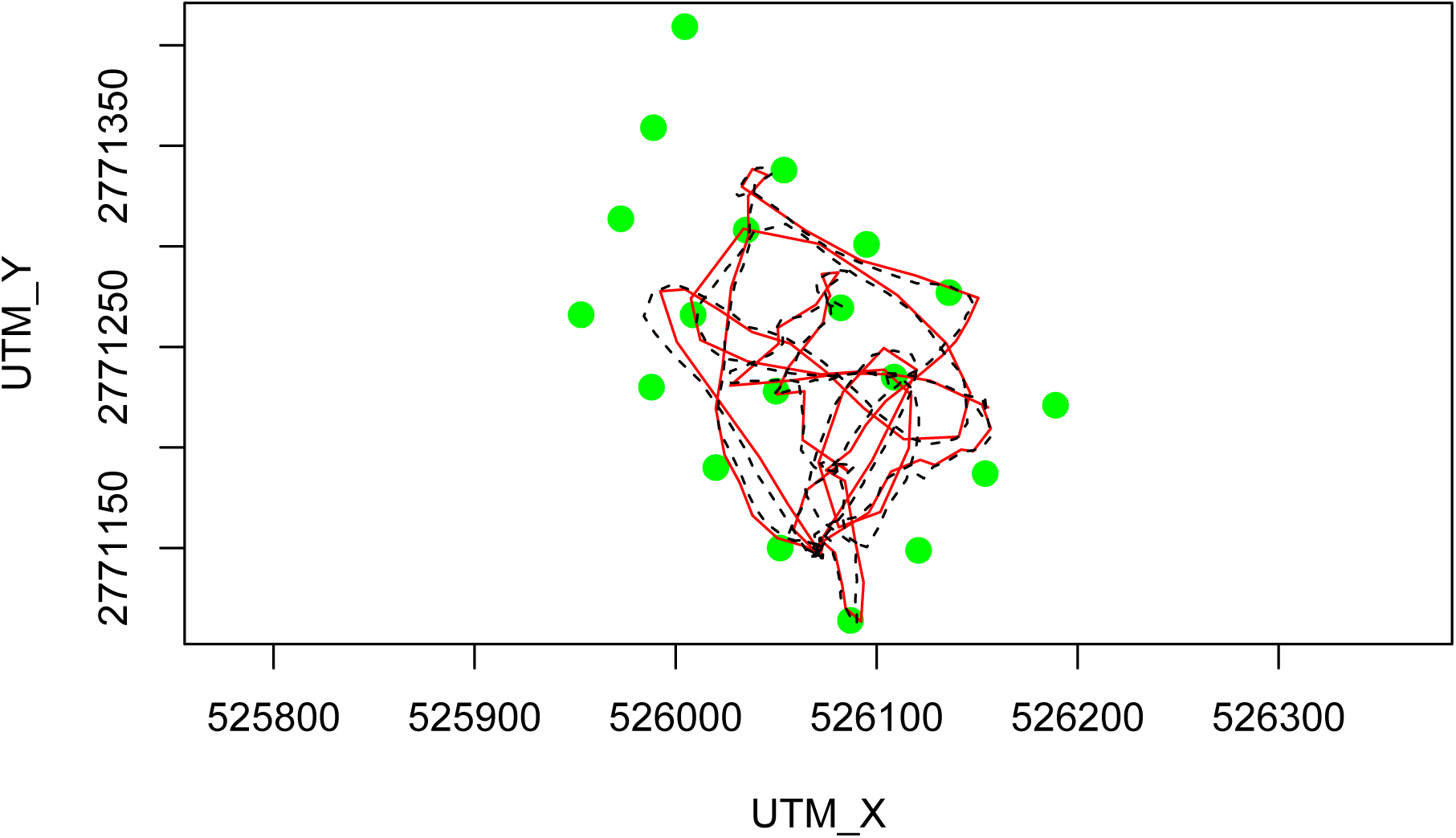
Estimated track (red line) and true track obtained using gps (black line). Green dots indicate hydrophone positions.

Basic code to plot X and Y coordinates separately.

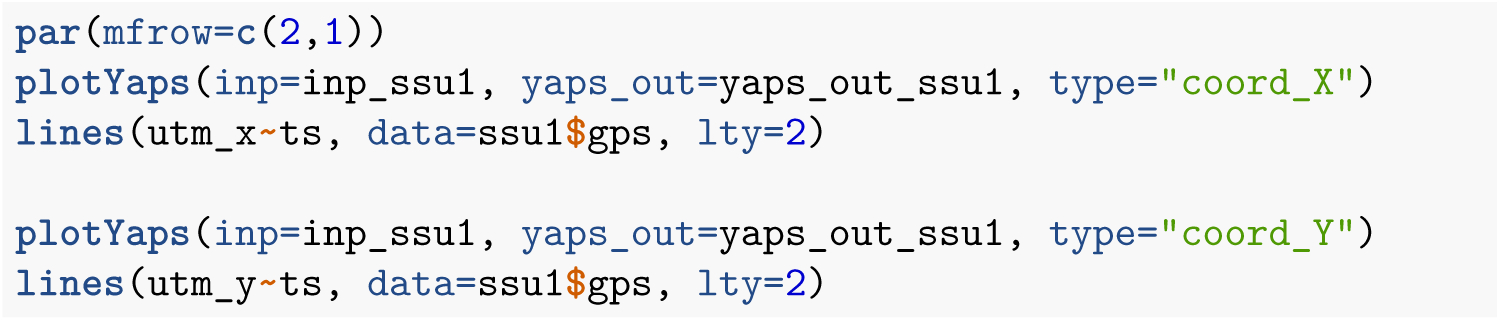

**Figure 9:**
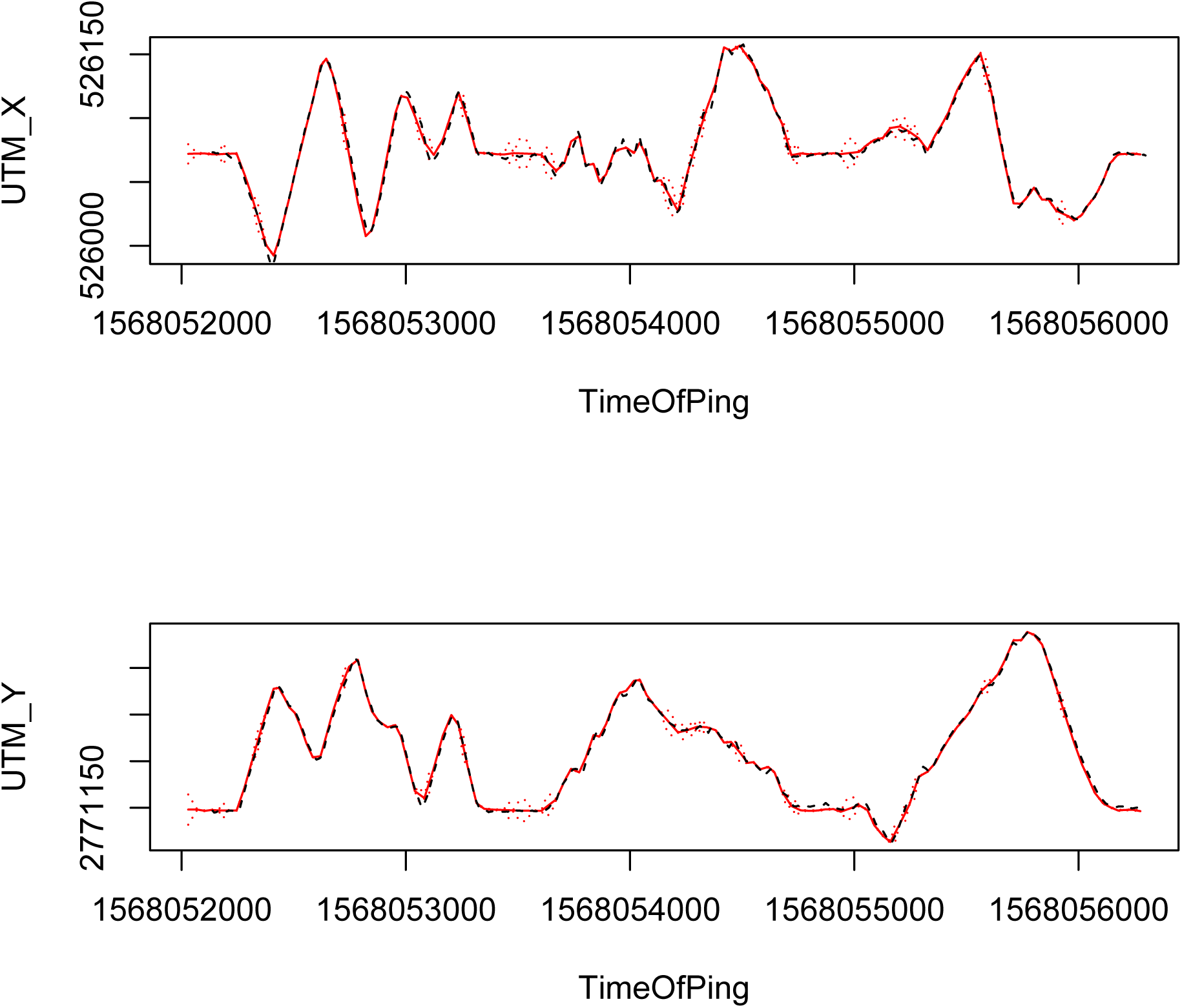
X and Y coordinates of true track (black) and estimated track (red). Broken red lines indicate estimated track +-standard error of position estimate.

## Discussion

Estimating positions and tracks of acoustically tagged fish based on data collected by autonomous receivers in a variable and often noisy environment is not a trivial task. Many factors, such as acoustic environment (e.g. degree of echoing from hard substrates and ambient noise level), water temperature and user experience level can influence both data quality and the level of pre-processing needed. Additionally, hydrophone performance and output format vary considerably between (and even within) manufacturers. Furthermore, hydrophones do not always behave and perform as expected. For instance, some hydrophone models autonomously initiate reboots causing perturbation of varying magnitude and/or duration of the internal clock at apparently random time intervals. Therefore, the functions in `yaps` might perform sub-optimal or fail miserably when applied to new data, if all these factors are not accounted for correctly. To that end, we want to emphasize that quality of `yaps` output is highly depended on quality of the input. Therefore, users should not expect `yaps` to salvage a poorly designed study or bad quality detection data.

